# Hybrid assembly of the large and highly repetitive genome of *Aegilops tauschii*, a progenitor of bread wheat, with the mega-reads algorithm

**DOI:** 10.1101/066100

**Authors:** Aleksey V. Zimin, Daniela Puiu, Ming-Cheng Luo, Tingting Zhu, Sergey Koren, James A. Yorke, Jan Dvorak, Steven L. Salzberg

## Abstract

Long sequencing reads generated by single-molecule sequencing technology offer the possibility of dramatically improving the contiguity of genome assemblies. The biggest challenge today is that long reads have relatively high error rates, currently around 15%. The high error rates make it difficult to use this data alone, particularly with highly repetitive plant genomes. Errors in the raw data can lead to insertion or deletion errors (indels) in the consensus genome sequence, which in turn create significant problems for downstream analysis; for example, a single indel may shift the reading frame and incorrectly truncate a protein sequence. Here we describe an algorithm that solves the high error rate problem by combining long, high-error reads with shorter but much more accurate Illumina sequencing reads, whose error rates average <1%. Our hybrid assembly algorithm combines these two types of reads to construct *mega-reads*, which are both long and accurate, and then assembles the mega-reads using the CABOG assembler, which was designed for long reads. We apply this technique to a large data set of Illumina and PacBio sequences from the species *Aegilops tauschii*, a large and highly repetitive plant genome that has resisted previous attempts at assembly. We show that the resulting assembled contigs are far larger than in any previous assembly, with an N50 contig size of 486,807. We compare the contigs to independently produced optical maps to evaluate their large-scale accuracy, and to a set of high-quality bacterial artificial chromosome (BAC)-based assemblies to evaluate base-level accuracy.

## Introduction

Long-read sequencing technologies have made significant advances in the past few years, with read lengths rapidly increasing while costs steadily dropped. Current technology can yield reads with average lengths of 5-10 kilobases (Kb) and a throughput that can reach a gigabase (Gb) from a single Pacific Biosciences (PacBio) SMRT cell. Although this technology remains more expensive and has lower throughput than Illumina short-read sequencing, it is now feasible to generate deep coverage of a large plant or animal genome at a modest cost. The long read lengths are extremely valuable for *de novo* genome assembly, allowing assemblers to overcome many of the problems caused by repeated sequences. This is particularly true for plant genomes in which transposable elements with lengths greater than 1 Kb are pervasive, often occupying over half of the genome. In the absence of other linking information, any near-exact repeat longer than a read will create a break in an assembly.

Traditionally, the primary strategy for spanning long repeats has been to create paired-end libraries from long DNA fragments, ranging in size from 2-10 Kb, or from even longer fosmids (˜40 Kb) or BACs (˜125-150 Kb). These strategies yield valuable long-range linking information, but they require more complex and more expensive methods of preparing DNA so that both ends can be sequenced. In contrast, when a single read spans a repeat and contains unique flanking sequences, the repeat can be directly incorporated into the assembly without the need to use paired-end information.

Recently, several assembly techniques have been developed for *de novo* assembly of a large genome from high-coverage (50x or greater) PacBio reads. These include the PBcR assembler, which employs the MHAP algorithm (Berlin et al. 2015) together with the CABOG assembly system; the HGAP assembler (Chin et al. 2013), the CANU assembler (https://github.com/marbl/canu) which also uses MHAP; and the Falcon assembler developed at Pacific Biosciences (https://github.com/PacificBiosciences/FALCON). Other methods employ a hybrid assembly strategy, in which short Illumina reads are used to correct errors in longer PacBio reads (Koren et al. 2012; Hackl et al. 2014; Salmela and Rivals 2014).

In this paper we describe a new hybrid assembly technique that can produce highly contiguous assemblies of large genomes using a combination of PacBio and Illumina reads. The new method extends Illumina reads into super-reads (Zimin et al. 2013) and then combines these with the PacBio data to create *mega-reads*, essentially converting each PacBio read into one or more very long, highly accurate reads. The mega-reads software, which is now incorporated into the MaSuRCA assembler, can handle hybrid assemblies of almost any plant or animal genome, including genomes as large as the 22 Gbp loblolly pine. We use this method to produce an assembly of the large and complex genome of *Aegilops tauschii*, one of the three diploid progenitors of bread wheat. The *Ae. tauschii* genome is unusually repetitive and has proven extremely difficult to assemble using short read data.

*Ae. tauschii* is a cleistogamic inbreeder, and its genome is nearly homozygous, making it an excellent asset for evaluation of error rates in assembly. To this end, we have generated an optical BioNano genome (BNG) map for the *Ae. tauschii* genome, which provided a sequence-independent means of evaluating the large-scale accuracy of the assembly, as we discuss below. We have also independently sequenced the *Ae. tasuchii* genome using an ordered BAC-clone sequencing approach, which provides a means of evaluating the base-level accuracy of the assembly.

## Computational Methods

### Sequencing data requirements

Our assembly recipe calls for at least 100x genome coverage by paired Illumina reads of 100-250bp, combined with at least 10x coverage in PacBio reads. Based on preliminary data, we expect that generating deeper PacBio coverage, up to 60x, is likely to improve the final results. The mega-reads algorithm has the following main steps.

### Super-read construction

We first transform Illumina paired-end reads into *super-reads*, as described previously (Zimin et al. 2013). The super-reads algorithm builds a database of all sequences of a user-specified length k, and then extends these k-mers in both directions as long as the extensions are unambiguous. In most cases, the super-reads will be much longer than the original Illumina reads, typically averaging 400 bp or more, depending on the repetitiveness of the genome. Subsequent steps of our algorithm use the longer but much lower coverage (usually 2-4x) super-reads, thus providing a very substantial degree of data compression. Another benefit of the longer super-reads is that they can be mapped to the high-error PacBio reads much more reliably than the shorter Illumina reads, and therefore provide a better vehicle for error correction.

For the next two steps, we treat each PacBio read as a template to which super-reads can be attached, as illustrated in **Figure 1**.

**Figure 1.**
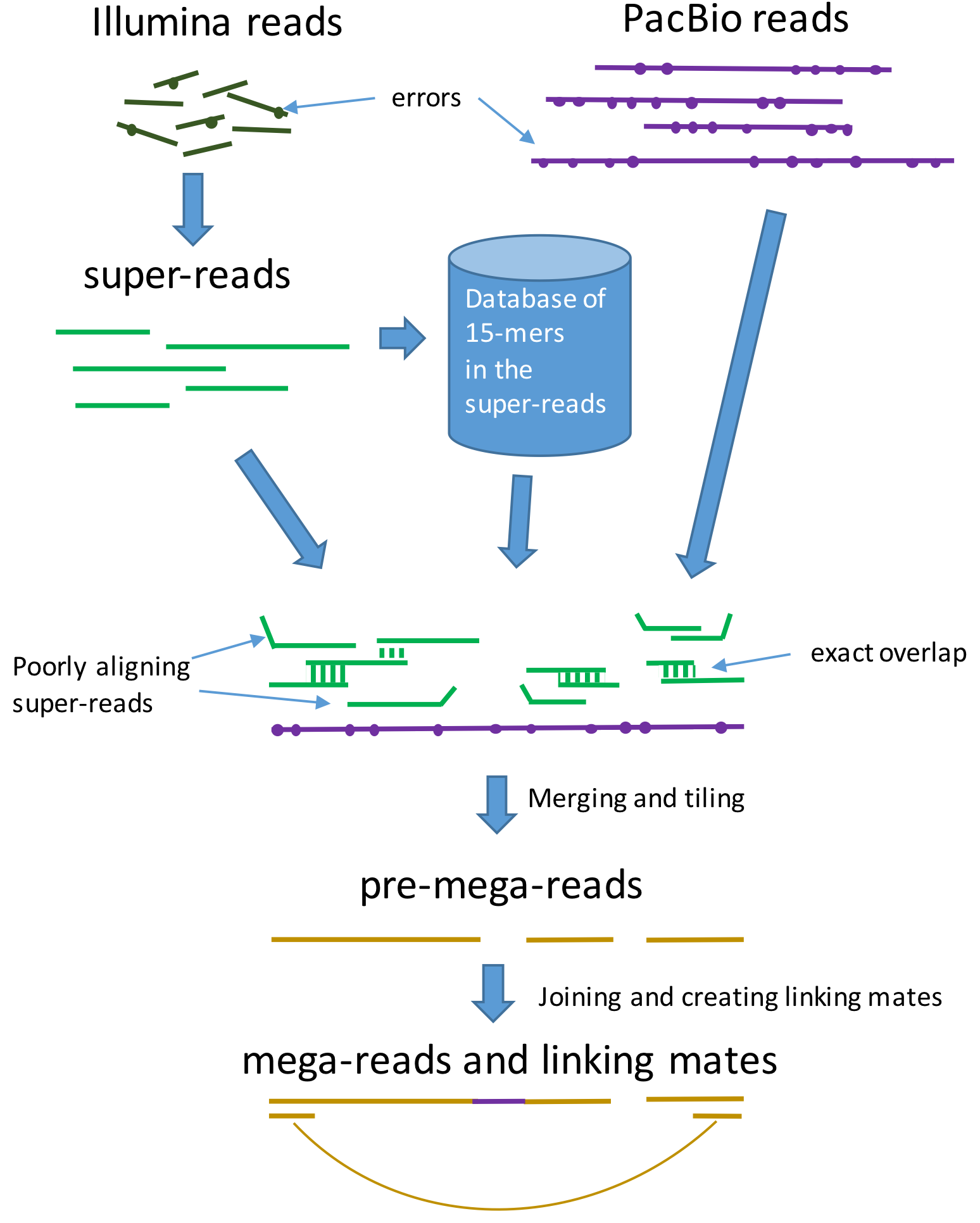
Overview of the mega-reads algorithm. Low-error rate Illumina reads (top left) are used to build longer super-reads (green lines), which in turn are used to construct a database of all 15-mers in those reads. PacBio reads (purple lines) and super-reads are then aligned, using the 15-mer database. Inconsistent super-reads are show as kinked lines; these are discarded and the remaining super-reads are merged, using the PacBio read as a template, to produce pre-mega-reads (yellow). These are further merged to produce the final mega-reads and to generate linking mates across gaps.

### Approximate alignment along a PacBio read

We create approximate alignments of the super-reads to each PacBio read using 15-mers that the PacBio read has in common with super-reads. (Note that the choice of k=15 for this step was made empirically, but it can be changed for different data.) We first build a database of all 15-mers in the super-reads, and use this database to compute, for each super-read, its approximate start and end positions on each PacBio read. This approach is similar (although different in many details) to both MHAP (Berlin et al. 2015) and minimap (Li 2016), in that both these other algorithms find chains of “seed” alignments in long PacBio reads. Our method does not compute a full alignment.

For each PacBio read *P*, we walk down the read looking at each 15-mer. We use the 15-mer database to determine (in constant time for each 15-mer) which 15-mers are found in super-reads. Once we have the super-reads that match *P*, for each such super-read *S* we look for ordered subsequences of the 15-mers that both *P* and *S* have in common. (The 15-mers can be overlapping.) We then assign a score to each super-read S, where the score is number of 15-mers in the longest common subsequence (LCS) of 15-mers in the two reads. We label an alignment as **plausible** if the score of *S* exceeds some specified minimum. For each plausible alignment, we compute an approximate position of *S* along *P* based on the positions of the LCS 15-mers in *P* and *S*.

Note that a super-read can align to many different PacBio reads; the number will depend on the depth of coverage of the PacBio data.

### Graph traversal for a PacBio read

Let K denote the K-mer size that was used to generate the super-reads. After super-reads are constructed, we record all exact overlaps of pairs of super-reads for which the length of the overlap is at least K.

Using all super-read positions on a PacBio read *P*, we create possible paths of (plausible) super-reads along *P*. Each path consists of a sequence of super-reads where two adjacent super-reads must have an exact overlap of at least K bases, and also must have positions on *P* that make it possible for them to overlap. We compute an LCS score for each path.

A path might span only part of *P*, and conversely subsequences of *P* might not be covered by any path. We then form a graph consistiing of the paths along *P*, where super-reads are the nodes and K-overlaps are the edges. For each connected component of paths (or more precisely a connected graph of super-reads), we compute the LCS score and we choose the path with the highest score. We call each such path a *pre-mega-read*. The sequence of a mega-read is essentially a long, high-quality “read” that covers part or all of the original PacBio read *P*.

At this point, each connected component is a directed acyclic graph (DAG) of super-reads that overlap by at least K bases and that align to *P*. The approximate positions of the super-reads on *P* impose a topological order on the DAG. We impose an overall direction on the DAG from the 5’ end towards the 3’ end of *P*.

### Tiling

We tile the PacBio read *P* with the pre-mega-reads in a greedy fashion, beginning with the longest pre-mega-read, and disallowing overlaps longer than K bases (**Figure 1**). We choose the pre-mega-reads for *P* by maximizing the total of all LCS scores in the tiling.

Many PacBio reads will be tiled by more than one pre-mega-read; i.e., the tiling has gaps. Gaps might be caused by lack of Illumina read coverage for parts of the genome, or by long stretches of poor quality sequence in a PacBio read, or (rarely) by chimeric PacBio reads. Even though we have PacBio sequence spanning these gaps, we choose not to simply merge the pre-mega-reads using raw PacBio read sequence because that might create stretches of low-quality sequence in the mega-reads. However, if multiple PacBio reads overlap one another for the sequence in one of these gaps, we can sometimes fill the gap between pre-mega-reads. We only use raw PacBio read sequence if 3 or more PacBio reads have nearly identical gaps, for which the pre-mega-reads surrounding the gap are identical and the gap lengths are nearly identical. In these cases, we create a consensus sequence from the multiple PacBio reads and use that to fill the gaps in each of them.

It is also possible that a gap in the tiling is not a gap at all, but instead is an erroneous insertion in the PacBio read. In these cases, the pre-mega-reads flanking the gap may overlap one another. If the pre-mega-reads overlap by at least 37 bp, then we merge them to close the gap.

Note that the user can set the maximum gap size for the gap-filling procedure, and the algorithm will not attempt to fill gaps larger than this maximum. If one sets the maximum gap size to zero, the megareads assembler will not use raw PacBio sequence at all, and will only join pre-mega-reads when they have an exact overlap of 37 bases or more.

The result of this tiling and gap-filling process is the final set of mega-reads.

### Creating linking pairs

When mega-reads cannot be merged and a gap remains, we create a linked pair of “reads” that spans the gap. We extract two 500 bp sequences from the mega-reads flanking the gap and link them together as mates (**Figure 1**). (If either mega-read is <500 bp, we create a shorter linking read.) The assembler uses these sequences in its scaffolding step to ensure that all mega-reads from the same PacBio read are kept adjacent in the assembly; i.e., they are placed into the same scaffold. We call these artificial mates the linking pairs.

### Assembly

Finally, we assemble the mega-reads along with the linking pairs into contigs and scaffolds using the CABOG assembler (Miller et al. 2008). For this step we can also use other linking information, if available, for scaffolding.

## Results

### Data sets

We generated over 19 million PacBio reads, equivalent to ˜38x genome coverage, using the SMRT P6-C4 chemistry. We also generated a total of 177x coverage on an Illumina HiSeq 2500 in paired 200 bp reads and an Illumina MiSeq with paired 250 bp reads (**Table 1**). These data sets were the only input used for our hybrid assembly of *Ae. tauschii*.

**Table 1.** Input data used for the *Ae. tauschii* hybrid assembly. Coverage is computed based on an estimated genome size of 4.25 Gb. Paired Illumina reads were generated from fragments whose lengths averaged 450-500 bp.

|  |  |  |  |
| --- | --- | --- | --- |
| Sequence data type | Number of reads | Average read length (bp) | Genome coverage |
| Illumina HiSeq paired-end | $1.98 \times 10^9$ | 200 | 93.2x |
| Illumina MiSeq paired-end | $1.41 \times 10^9$ | 250 | 83.6x |
| PacBio SMRT P6-C4 | $19.22 \times 10^6$ | 8,519 | 38.5x |

### *Ae. tauschii* assembly

The methods described above produced 16.7M super-reads from the Illumina data, and 18.7M mega-reads from the super-reads and PacBio reads (**Table 2**). We aligned both superreads and mega-reads to an Illumina-only assembly, produced by the DeNovoMagic assembler, and the average identity was 99.91% and 99.77% respectively. From this we estimate that the error rates for super-reads and mega-reads were less than 0.09% and 0.23% (respectively) because some of the mismatches might be caused by haplotype differences or by errors in the Illumina-only assembly.

**Table 2.** Statistics for super-reads and mega-reads. Super-reads were constructed from Illumina data, and mega-reads were constructed as described in the main text. Coverage is based on an estimated genome size of 4.25 Gbp. Error rates were computed by mapping the reads against Illumina-only contigs.

|  |  |  |  |  |  |
| --- | --- | --- | --- | --- | --- |
|  | Number | Coverage | Average length (bp) | N50 (bp) | Error rate (%) |
| Super-reads | $16.7 \times 10^6$ | 1.9x | 474 | 749 | <0.09 |
| Mega-reads | $18.7 \times 10^6$ | 27.8x | 6,319 | 9,378 | <0.23 |

We next ran the CABOG assembler (version wgs-8.3rc2) to produce an initial assembly of the mega-reads. This assembly contained 128,898 scaffolds totaling 4.778 Gb in length. We then ran four rounds of alignment, aligning each scaffold to all others, to remove scaffolds that were either duplicated or that were completely contained within other scaffolds. For this alignment step we ran bwa-mem (Li 2013) with parameters `–k127 –e` and then used nucmer (Delcher et al. 2002a) to find and remove duplicate alignments. This procedure identified a total of 75,338 scaffolds (most of them very small) that were contained in other scaffolds and could be safely removed. Total computational time for all steps of the assembly was approximately 110,000 CPU hours, with about 72,000 CPU hours for computing superreads and mega-reads, and the remaining time spent in assembly of contigs and scaffods. The code is highly parallelized so that most procedures were run in parallel on large computing grids.

The resulting *Ae. tauschii* assembly, version Aet_MR.1.0, contains 53,560 scaffolds with a total span of 4.338 Gb, a contig N50 size of 486,807 bp, and a scaffold N50 size of 521,653 bp (**Table 3**). As described above, scaffolding was minimal because it used only the linking pairs created from mega-reads that flanked gaps in the original PacBio reads. Thus for every gap internal to a scaffold, we have at least one PacBio read spanning the gap. The principal benefit of PacBio reads and of the mega-reads algorithm is the much larger contigs that result, ˜30 times larger than the contigs from an Illumina-only assembly (**Table 3**).

**Table 3.**
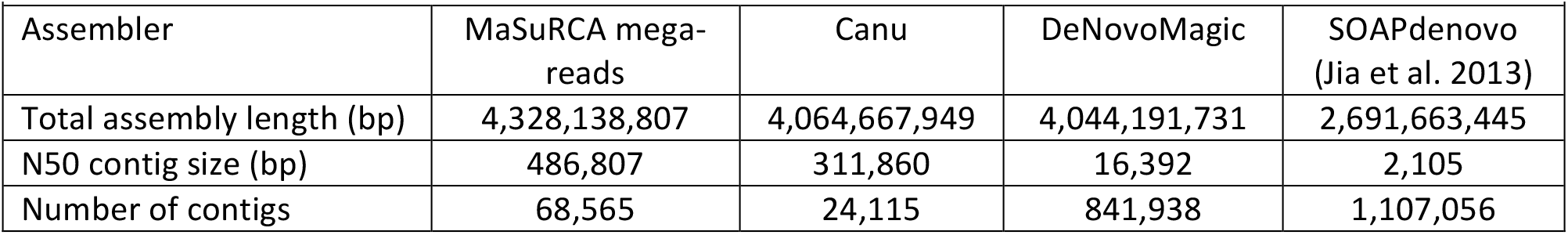
Assembly statistics for *Ae. tauschii* Aet_MR.1.0 compared to other assemblies. N50 numbers were computed using an estimated genome size of 4.25 Gb. The Canu assembly used PacBio data only but at deeper coverage, 55x. The DeNovoMagic assembly used additional Illumina paired reads from fragments ranging in size from 3-10 Kb. The SOAPdenovo assembly (Jia et al. 2013) used 94x coverage in Illumina and 454 sequences. The MaSuRCA mega-reads assembly used only the short-fragment Illumina pairs plus the PacBio data.

| Assembler | MaSuRCA mega-reads | Canu | DeNovoMagic | SOAPdenovo (Jia et al. 2013) |
| --- | --- | --- | --- | --- |
| Total assembly length (bp) | 4,328,138,807 | 4,064,667,949 | 4,044,191,731 | 2,691,663,445 |
| N50 contig size (bp) | 486,807 | 311,860 | 16,392 | 2,105 |
| Number of contigs | 68,565 | 24,115 | 841,938 | 1,107,056 |

We also created a whole-genome assembly with the PacBio reads only, using the Canu assembler (Berlin et al. 2015). For this assembly we generated additional data to bring the PacBio coverage up to 55x. The Canu assembly had a total length of 4.06 Gb and an N50 size (using 4.25 Gb as the genome size) of 311,860 bp (**Table 3**). It is worth noting here that the Aet_MR.1.0 assembly has many more contigs than the Canu assembly, but this is due to a large number of small contigs in the tail of the distribution, and to the fact that the Aet_MR.1.0 assembly is ˜264 Mb larger. If we select contigs from the Aet_MR.1.0 assembly whose sizes total 4.06 Gb (the same total as the Canu assembly), we need only 24,309 contigs, almost exactly the same number as in the Canu assembly, and the smallest such contig is 15,911 bp.

### Evaluation of assembly quality

We evaluated the quality of the output contigs using two metrics: large-scale contiguity and consensus sequence accuracy. To evaluate large-scale accuracy we aligned the Aet_MR.1.0 contigs to independently developed genome maps created using nanochannel arrays from BioNano Genomics. This technology allows the construction of accurate maps based on restriction enzymes, in which DNA molecules are passed through a nanochannel and fluorescently tagged restriction sites are detected (Lam et al. 2012). This process creates many small restriction-mapped regions that can span several megabases each. Nanochannel maps have recently been used to improve the assembly of portions of several highly repetitive plant genomes, including one arm of the bread wheat genome (Stankova et al. 2016), a 2-Mb fragment of *Ae. tauschii* (Hastie et al. 2013), and six small but complex regions of the maize genome (Dong et al. 2016). To use these maps to assess quality of a sequence assembly, the distances and positions of the same restriction sites along the nanochannel map (“nanomap”) contigs are compared with a restriction map constructed computationally.

Here we used the Ae. tauschii nanomap only to evaluate the correctness of Aet_MR.1.0; it was not used to construct or modify the assembly. We aligned our contigs to the Ae. tauschii nanomap using the restriction sites in the contigs, and searched for regions of disagreement, which indicate either an error in the nanomap or a mis-assembled contig. Note that the nanomap does not contain sequence, and the only errors this procedure can detect are relatively large-scale rearrangements, insertions, or deletions.

We found 572 locations where a contig disagreed with the nanomap. A typical signature of a possible mis-assembly is a sharp drop in coverage, indicating a possible “weak” overlap holding the contig together. Given that the average coverage in mega-reads was ˜28x (**Table 3**), we flagged as a possible mis-assembly any disagreement with the nanomap where the coverage was <4; this analysis flagged 342 locations. We examined a small set of the higher-coverage discrepancies by hand, and in each case the assembly appeared correct (based on underlying support from paired Illumina reads); thus we concluded that these are likely to represent errors in the nanomap rather than in Aet_MR.1.0. Thus for the full assembly of 4.28 Gb, we estimate approximately 1 assembly error per 12.5 Mb.

Next we evaluated the base-level sequence quality by aligning the contigs to 5,216 assembled BAC pools, each containing eight overlapping BACs with approximately 1 Mb per pool. These pools were independently assembled from 250-bp Illumina MiSeq reads using SOAPdenovo2 (Luo et al. 2012), and previously released as a preliminary assembly (Aet 0.4) at ftp://ftp.ccb.jhu.edu/pub/data/Aegilops_tauschii. The reads used for these BAC pool assemblies were a subset of reads we used for creating the mega-reads. We aligned all contigs to the BAC pool assemblies with Nucmer program from the MUMmer package (Delcher et al. 2002b; Kurtz et al. 2004) with a minimum match length of 127 bp to anchor each alignment.

We then computed the average identity for all alignments longer than 10 Kb, which is long enough to span almost all repeats in the genome. This yield 10,597 alignments covering 1,629,700,561 bp, and the average identity was 99.96%, with values ranging from 99.24% to 100%. Thus the base-level accuracy of the assembly appears very high: note that some differences between the haploid BAC assemblies and the diploid whole-genome assembly are likely due to haplotype differences. In support of this hypothesis, if we consider only the alignments at 99.99% identity, these cover 650,572,225 bp (40%) of the aligned regions.

## Discussion

Both *Ae. tauschii* and its close relative, the hexaploid wheat *(Triticum aestivum)*, have proven difficult to assemble because of their unusually high proportion of repetitive sequences. A previously published version of *Ae. tauschii* (Jia et al. 2013) yielded only 2.69 Gbp (˜63% of the genome) spread across 1.1 million contigs. Attempts to assemble *T. aestivum* have met with similar problems: a massive effort to sequence *T. aestivum* chromosome-by-chromosome yielded only 61% of the genome in very small contigs with N50 sizes from 1.7 to 8.9 Kb (International Wheat Genome Sequencing 2014). Most of the repeats in *Ae. tauschii* and in other plants consist of transposons (Lisch 2013), which occur in thousands of copies, many of them nearly identical, throughout the genome. Assembly algorithms can find the correct location for these elements if the input data include reads that are long enough to contain the entire span of a repeat plus unique flanking regions on either side.

The PacBio reads generated in this study, with an average read length of 8520 bp, are easily long enough to span most transposable elements, which are usually 2-3 kilobases in length (though some can be longer). However, the high error rate of PacBio reads requires some form of error correction before these sequences can be used in a final assembly. The mega-reads introduced here solve both these problems: with an average length of 6319 bp, they are long enough to contain the ubiquitous 2-3 Kbp repeats in the *Ae. tauschii* genome, and they are accurate enough–much more accurate than raw Illumina reads–to be used to generate a high-quality assembly. Using these mega-reads, we have generated a whole-genome assembly of *Ae. tauschii* with an N50 contig size of 486,807 bp, more than 20 times longer than any previous assembly. The unprecedented contiguity of this assembly provides a strong foundation for additional mapping and assembly work to create a far more complete picture of this important plant genome. The strategy described here, using deep coverage Illumina sequencing with moderate coverage PacBio sequencing, demonstrates a cost-effective approach to generating highly contiguous, accurate assemblies of large genomes, even when those genomes contain large numbers of long, near-identical repeats.

### Data and software availability

The MaSuRCA mega-reads software is freely available from http://genome.umd.edu/masurca.html. The *Ae. tauschii* assembly, version Aet_MR.1.0, is available from NCBI under BioProject PRJNA329335.

## Acknowledgements

This work was supported in part by NSF grant IOS-1238231 to J.D. and by NIH grant R01 HG006677 to S.L.S. S.K. was supported by the Intramural Research Program of the National Human Genome Research Institute, NIH. This study utilized the computational resources of the Biowulf system (http://biowulf.nih.gov) at NIH and the MARCC system (http://marcc.jhu.edu) at JHU.

